# *sabinaHSBM*: An R package for link prediction network reconstruction using Hierarchical Stochastic Block Models

**DOI:** 10.1101/2025.10.28.684773

**Authors:** Herlander Lima, Jennifer Morales-Barbero, Rubén G. Mateo, Ignacio Morales-Castilla, Miguel A. Rodríguez

**Author notes:** Equal contribution (Herlander Lima, Jennifer Morales-Barbero).

## Abstract

1. Network analysis is a powerful framework for investigating complex systems across disciplines, including ecology, where it helps uncover patterns in predator–prey, host– parasite, or plant–pollinator interactions. However, ecological network data are often incomplete or error-prone due to sampling limitations, detection failures, and taxonomic uncertainty—leading to missing (false negative) and spurious (false positive) links that obscure structure and hinder inference. The hierarchical stochastic block model (HSBM), particularly in its degree-corrected form, is among the most effective tools for reconstructing networks under such uncertainty. Despite its robustness, the primary implementation of HSBM in the Python-based graph-tool library has remained largely inaccessible to ecologists.
2. Here, we introduce *sabinaHSBM*, the first R package that makes degree-corrected HSBM broadly available through a user-friendly, flexible workflow. By bridging a gap between advanced network modeling and widely used ecological analysis platforms, *sabinaHSBM* facilitates network reconstruction and link prediction from binary bipartite data. The workflow involves three main steps: (1) preparing input data, (2) estimating posterior link probabilities, and (3) reconstructing the network. The package supports detection of undocumented and spurious links, exploration of hierarchical structure, and propagation of uncertainty throughout. Key features include cross-validation, flexible thresholding, probabilistic evaluation metrics, and two link prediction modes: estimating all link probabilities or identifying undocumented ones.
3. We illustrate the package’s functionality through a case study using a published global dataset of carnivore–parasite associations, showing that inferred groupings are phylogenetically clustered. To assess predictive accuracy, we examined the top 10 highest-probability links identified by the model and found published evidence for 8, despite their absence from the original dataset. This highlights the model’s ability to recover biologically meaningful but underreported interactions.
4. By integrating all components of HSBM-based reconstruction into an accessible R package, *sabinaHSBM* empowers researchers to improve relational data quality and uncover overlooked patterns in complex ecological networks and beyond.

## 1 INTRODUCTION

Network analysis, widely used across disciplines from neuroscience and epidemiology to social sciences and climate research, provides a powerful framework for representing and analyzing complex systems. In ecology, it has become central to exploring species interactions through bipartite networks, which represent relationships between two sets of entities—such as predators and prey, hosts and parasites, or plants and pollinators—as nodes connected by edges denoting observed links. These networks are essential tools for understanding biodiversity patterns, coevolutionary dynamics, and ecosystem functioning (Delmas et al., 2019). However, empirical ecological data are almost invariably incomplete, noisy, and biased. Sampling limitations, detection errors, accessibility issues, and taxonomic ambiguities introduce false negatives (missing links) and false positives (spurious links), obscuring true network structure and compromising the robustness of ecological inference (Delmas et al., 2019; Fründ et al., 2016; Morales-Castilla et al., 2015). Treating such data as static, error-free adjacency matrices risks misrepresenting the systems they describe and may limit their usefulness in research and data-informed conservation efforts.

Stochastic block models (SBMs) are foundational tools in network science (Holland et al., 1983), providing a flexible probabilistic framework for detecting latent community structure in network data. Among the most effective extensions for recovering such structure and addressing uncertainty in bipartite networks is the degree-corrected hierarchical stochastic block model (HSBM), developed by Peixoto (2014, 2018), which underpins the R package we introduce here: *sabinaHSBM* (see below). This formulation explicitly accounts for variation in node degree within blocks—a common feature of real-world bipartite datasets—thereby reducing misclassification caused by uneven sampling. By treating the observed adjacency matrix (i.e., the *n* × *n* matrix with 1/0 entries indicating whether each node pair is observed or not) as a noisy realization of an underlying generative process, HSBM can uncover both community structure and latent hierarchical patterns (T. P. Peixoto, 2018), effectively assigning probabilities to both documented and undocumented links. Indeed, in a recent benchmark involving over 200 algorithms and 550 empirical networks from six scientific domains, HSBM was identified as the most accurate individual method for link prediction (Ghasemian et al., 2020). Similarly, in a focused comparison of ecological networks, HSBM outperformed alternative approaches (Terry & Lewis, 2020).

Beyond its strong predictive performance, HSBM also offers several methodological advantages over alternative network reconstruction methods. First, as a nonparametric Bayesian framework, it can infer hierarchically nested group structures and estimates posterior link probabilities based solely on network topology; that is, without relying on predefined structural constraints or parametric assumptions about link distributions. Although it is theoretically grounded in a generative model based on SBM (Holland et al., 1983), the framework flexibly adapts to the data, effectively relaxing this assumption in practice. Moreover, unlike many statistical or machine learning methods, HSBM does not require external covariates or metadata, making it especially well-suited for ecological datasets where such information is often sparse or uncertain—and where meaningful structure must be inferred directly from the network itself. For instance, users are not required to define the number of communities (also known as modules, groups or blocks), as these are automatically inferred from the observed network. Second, it enables network reconstruction from a single measurement, even without prior error estimations. This capability addresses a major limitation in observational datasets, where repeated measurements are often impractical or unavailable. Third, HSBM quantifies uncertainty by generating posterior edge probabilities, enabling researchers to assess link reliability. This approach can serve to support the identification of both reliable and uncertain links, facilitate informed decision-making, prioritize sampling efforts, and refine or validate existing hypotheses. Finally, HSBM leverages the inherent structure of networks to detect hierarchical groupings or communities, identifying the structural organization of the network. This output provides insights into the network’s architecture at multiple levels of resolution, uncovering patterns that may be otherwise overlooked, and offering a valuable interpretation tool beyond standard flat clustering.

Despite the notable characteristics of the HSBM, its original implementation relies on the *graph-tool* module (Peixoto, 2014; 2018), a Python package that operates exclusively on Unix systems. This limitation hinders its accessibility to researchers who primarily work with R programming and Windows systems. The *sabinaHSBM* addresses this gap by providing a more accessible network reconstruction tool tailored for R users.

In the following sections, we describe the design and implementation of *sabinaHSBM* (Section 2), demonstrate its application to a mammal–parasite interaction network (Section 3), and discuss its broader methodological and ecological implications (Section 4).

## 2 THE *sabinaHSBM* PACKAGE

We introduce *sabinaHSBM*, an R package that provides a straightforward and user-friendly workflow for network reconstruction in three steps: (1) preparing the input database; (2) predicting link probabilities; and (3) reconstructing the network (Box 1). Each step is supported by dedicated functions (Table 1) that ensure flexibility, ease of use, and a wide range of outputs derived from the underlying HSBM engine.

**Table 1.**
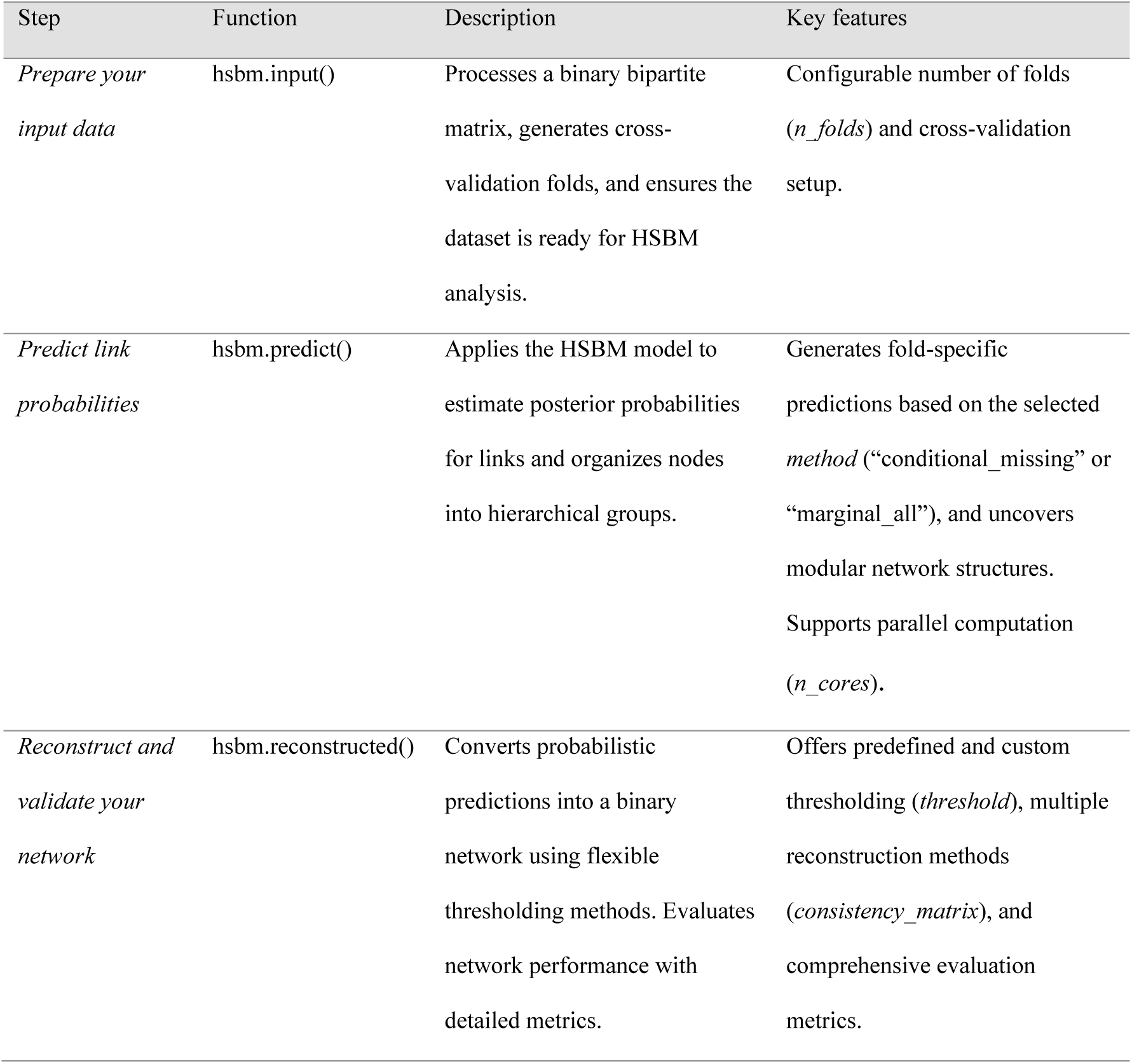
Overview of the *sabinaHSBM* workflow steps, associated functions, and their key features.

| Step | Function | Description | Key features |
| --- | --- | --- | --- |
| *Prepare your input data* | hsbm.input() | Processes a binary bipartite matrix, generates cross-validation folds, and ensures the dataset is ready for HSBM analysis. | Configurable number of folds (*n_folds*) and cross-validation setup. |
| *Predict link probabilities* | hsbm.predict() | Applies the HSBM model to estimate posterior probabilities for links and organizes nodes into hierarchical groups. | Generates fold-specific predictions based on the selected *method* (“conditional_missing” or “marginal_all”), and uncovers modular network structures. Supports parallel computation (*n_cores*). |
| *Reconstruct and validate your network* | hsbm.reconstructed() | Converts probabilistic predictions into a binary network using flexible thresholding methods. Evaluates network performance with detailed metrics. | Offers predefined and custom thresholding (*threshold*), multiple reconstruction methods (*consistency_matrix*), and comprehensive evaluation metrics. |

Designed to reconstruct bipartite binary networks from incomplete and biased data—a common limitation in ecological datasets—*sabinaHSBM* offers unique features. Built-in cross-validation tests generate held-out links across folds, ensuring that performance metrics reflect true generalizability. Held-out links are observed links (1s) that are treated as absences (0s) during model training to evaluate the model’s ability to recover them. Uncertainty quantification assigns each candidate interaction—i.e., link—a continuous posterior probability, enabling users to assess its reliability directly rather than relying on a simple matrix of predicted presences/absences. Our implementation of the package supports two prediction modes that serve different purposes: The “conditional_missing” mode implements a classical gap-filling strategy, computing conditional probabilities only for unobserved node pairs—ideal for conservative detection of missing interactions without questioning observed ones. The “marginal_all” mode embraces an innovative whole-network strategy (Peixoto, 2018). It repeatedly generates true latent graphs from the observed network and derives marginal probabilities for all observed and unobserved links—thereby uncovering both missing (false negatives) and spurious interactions (false positives) in one comprehensive analysis (see Section 2.2 for full details). Once the probabilities are computed, the package also provides flexible thresholding methods, different evaluation statistics—such as the area under the receiver operating characteristic curve (AUC, Fielding & Bell, 1997), the area under the precision-recall curve (AUC-PR) (Saito & Rehmsmeier, 2015), or the recovery link ratio (RLR, a name we propose here for the proportion of held-out links correctly recovered), among others—and adaptable consolidation approaches for generating the final reconstructed binary matrix/network, allowing researchers to adapt reconstructions to their objectives. Because HSBM inference is computationally demanding—especially across multiple cross-validation folds and/or large networks—*sabinaHSBM* integrates parallel processing via the *parallel* R package (R Core Team., 2024), distributing tasks across multiple cores for efficient execution. By leveraging the computational power and core functionalities for network reconstruction provided by *graph-tool* via the *reticulate* package (Ushey et al., 2025), *sabinaHSBM* enables researchers to execute Python-based computations seamlessly within R.

The current release can be installed directly from the GitHub repository with remotes::install_github(“h-lima/sabinaHSBM”).

Because *graph-tool* (the core engine) runs on Python/Unix, a ready-to-use Docker image is also available (https://hub.docker.com/r/herlima/sabinahsbm) to guarantee full functionality and accessibility on Windows, Mac, and Linux. That image packages the *sabinaHSBM* code, its R and Python runtimes (including *graph-tool*), all required libraries, and system dependencies into a single, portable container. Users simply pull this image and—following the guidance in Supporting Information S1—launch a container equipped with RStudio Server and Jupyter Notebook and all needed tools to run *sabinaHSBM*. This dual approach—reticulate integration and Docker compatibility—reduces technical barriers, extending the HSBM’s application to a broader audience and research contexts.

Supporting Information includes two end-to-end tutorials: Supporting Information S2 shows the “conditional_missing” pipeline for conservative missing link recovery, and Supporting Information S3 walks through the “marginal_all” pipeline for finding both missing and/or spurious links. Installation steps are provided at the beginning of each corresponding guide.

### 2.1 Step 1: *Prepare your input data with hsbm.input()*

The *sabinaHSBM* workflow begins by structuring input data for HSBM analyses using the hsbm.input() function. This function processes an observed binary bipartite matrix that represents interactions between two distinct sets of nodes (e.g., hosts and parasites in our example case study, but could be other interaction networks or distributional networks where nodes represent sites and species occurrences). Rows and columns correspond to these node types, and entries denote the presence (1) or absence (0) of a link. Any rows or columns without observed interactions are automatically removed. A summary() function for hsbm.input() objects provides a concise overview of matrix dimensions and observed links.

A key feature of hsbm.input() is the generation of cross-validation folds, which partition held-out links across groups. Users can customize the number of folds (*n_folds*, default: 5) or provide precomputed folds. In each fold, all observed links are included, but a different, randomly selected, non-overlapping subset of approximately equal size is masked as held-out, ensuring every observed link is excluded once during the cross-validation process. While setting *n_folds* = 1 is technically possible (used in combination with *no_heldout* = TRUE), it is not recommended, as it disables cross-validation and prevents the computation of classification thresholds for binary network reconstruction. However, it can be appropriate in post-validation stages to fit the model on the full network and generate final predictions. Although this configuration does not allow for threshold calculation, it enables ranking nodes by their probabilities based on the complete data structure. Further customization includes the minimum number of observations required per row and column (*min_per_row*, *min_per_col*, default: 2). The output includes the cleaned observed binary matrix (indicating species interactions in the example), cross-validation folds, and edgelists (i.e., tables listing node pairs) with metadata, such as edge types (e.g., documented or held-out links).

### 2.2 Step 2: Predict link probabilities with hsbm.predict()

In the second step, the function hsbm.predict() applies the HSBM model to identify hierarchical groupings of nodes (i.e., clusters of species or entities that tend to interact similarly) and generate probabilistic predictions for link existence. This function relies on the structured input from hsbm.input(), using the cross-validation folds to explore the internal structure of the network and to estimate the likelihood of interactions that may have been missed (false negatives) or wrongly recorded (false positives).

By default, predictions are computed across all folds, although a specific fold can be selected by specifying its index with the *elist_i* argument. To speed up execution on large datasets or multiple folds, the hsbm.predict() function supports parallel computation via the *n_cores* argument (default: 1), enabling the distribution of computations across multiple cores. For the network reconstruction task, hsbm.predict() assumes that the observed network *D* represents the best available information about the true, unobserved network *A*. Each entry *D_ij_* = 1 indicates that an edge between node pairs (*i*, *j*) was observed, whereas *D_ij_* = 0 denotes edges that were unobserved or removed temporarily as held out during cross-validation. The network *D* is treated as a noisy measurement of the true network *A*, potentially containing errors such as false positives (erroneously recorded edges) and/or false negatives (missing true edges). Following Peixoto (2018), a measurement model was employed where each node pair (*i*, *j*) in *D* is associated with *n_ij_* distinct measurements, resulting in *x_ij_* observed links, such that *D_ij_* = (*n_ij_*, *x_ij_*). Although this model accommodates arbitrary non-negative integers *x_ij_* ≤ *n_ij_*, *sabinaHSBM* focuses on the common case of binary adjacency matrices, where only a single measurement per node pair is available or the details of the measurement process are unknown. In this setting, *n_ij_* = 1 is assumed for all pairs (*i*, *j*) and *x_ij_* ∊ {0, 1} corresponds directly to the observed values in the adjacency matrix *D*.

The goal is to infer the true network *A* and the latent community structure *b* from the observed data *D **=*** (*n*, *x*), by targeting the joint posterior distribution *P*(*A*, *b* | *D*) ∝ *P*(*D*|*A*)*P*(*A*, *b*), where *P(D|A)* models the measurement process, and *P(A, b)* represents the prior joint probability of the network and its structure. Since the model uses the stochastic block model (SBM) as the generative prior, this can be decomposed as *P*(*A*, *b*) = *P*(*A*|*b*)*P*(*b*), where *P(A|b)* is the probability of network *A* given the block partition *b,* and *P(b)* is the prior on the block partition itself. This Bayesian formulation, detailed in Peixoto (2018), integrates uncertainty in the measurement process with the structural correlations captured by the HSBM. To explore the posterior, Markov Chain Monte Carlo (MCMC) sampling is used to generate *k* configurations (*A^(k)^*, *b^(k)^*) from the joint posterior distribution.

After an initial MCMC equilibration phase or burn-in to ensure convergence—controlled by the *wait* argument (default: 1,000 iterations)—the model generates a set of posterior samples (defined by the *iter* argument, default: 10,000 iterations) to estimate edge probabilities *p_ij_* (for details on the MCMC sampling process see Appendix 1 at Supporting Information S4). An optional *rnd_seed* argument allows fixing the random seed for the HSBM inference to ensure reproducibility.

Two methods for link prediction are available in *sabinaHSBM*, selectable via the *method* argument:

- The “conditional_missing” method (conditional probability averaging)

A classical gap-filling module that focuses exclusively on scoring unobserved links (i.e., those with *D_ij_* = 0). Specifically, for each unobserved node pair (*i*, *j*), it computes the average conditional probability that a link exists, given the inferred block structure.

After the initial MCMC equilibration or burn-in (controlled with *wait* argument), a set of *N* posterior block partitions *b^(k)^* is obtained (controlled with *iter* argument). For each partition, the package calls the MeasuredBlockState.get_edges_prob() function in *graph-tool* (http://graph-tool.skewed.de/) to compute the conditional probability *P*(*A*_*ij*_ = *1*| *b*^(*k*)^) and then averages these values across all *N* samples to yield:

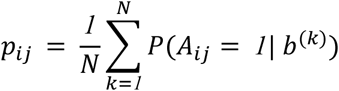

By relying exclusively on the inferred block structure, the “conditional_missing” method implements the classical approach for link prediction assigning each unobserved link a score based on its likelihood under a generative model. It retains the familiar/classical SBM link-prediction paradigm (e.g. Ghasemian et al., 2019; Aicher et al., 2015), allowing direct comparison with other existing algorithms. However, this method does not yield true marginal probabilities, and it is more computationally intensive than the “marginal_all” method. For these reasons, *sabinaHSBM* restricts this method to estimating probabilities for unobserved edges (*D_ij_* = 0) only.

- The “marginal_all” method (marginal posterior probability estimation)

A whole-network reconstruction approach that computes the marginal posterior probability *P*(*A*_*ij*_ = *1* | *D*) for observed (*D_ij_* = 1) and unobserved (*D_ij_* = 0) links. After the MCMC burn-in (controlled by the *wait* argument), we draw *N* samples (set via *iter* argument). For each sample we use the MeasuredBlockState.collect_marginal() function from *graph-tool* to generate a latent/sampled network *A^(k)^* and record every edge presence 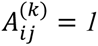. These edge presences are averaged to obtain:

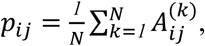

where *N* is the number of posterior samples and 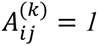 if the edge (*i*, *j*) exists in the *k*-th latent network, and 0 otherwise. The resulting *p_ij_* directly reflects the accumulated evidence for the existence of the edge (*i*, *j*) across the posterior. The “marginal_all” method delivers a marginal probability matrix that simultaneously uncovers missing links (false negatives) and spurious links (false positives), making it the method of choice for exhaustive network diagnostics and rigorous quality assessment.

Because it samples full networks rather than focusing on blocks, some links may never appear in any sample *A^(k)^*, yielding undefined *p_ij_*, *sabinaHSBM* lets you handle these cases flexibility via the *na_treatment* argument (see Section 2.3), ensuring no uncertainty is lost in downstream analyses.

Optionally, the function allows saving the results and *graph-tool* objects as Python pickle files. These files are stored in the working directory under names like hsbm_res_foldi.pkl, where *i* denotes the fold index. Also, a simple hierarchical edge bundling figure for each fold can be saved in the working directory, using the argument *save_plots* = TRUE.

The main outputs of the hsbm.predict() function are organized by fold and include the hierarchical group assignments revealing network modularity, available under $groups; minimum description length (MDL) values, found in $min_dl; and posterior probabilities *p_ij_* for links, based on the selected method for link prediction, accessible via the $probs attribute. These probabilistic output support downstream steps in the workflow, where users can reconstruct the network and evaluate the plausibility of individual interactions.

### 2.3 Step 3: Reconstruct and validate your network with hsbm.reconstructed()

Building on probabilistic predictions generated by hsbm.predict(), the hsbm.reconstructed() function generates evaluation metrics and a final binary network representation—i.e., a matrix of 1s and 0s indicating predicted observed/unobserved links—, which is essential for identifying missing and spurious links. It uses the *threshold* argument to transform link probabilities into these binary classifications, with several predefined thresholding methods available. These include the default “roc_youden”, which selects the threshold that maximizes the sum of sensitivity and specificity (Youden’s J statistic; Youden, 1950); “prc_closest_topright”, which minimizes the distance to the top-right corner of the precision-recall curve; or “prc_max_F1”, which optimizes the F1 score, among others. While the default threshold performs well in most cross-validation settings and is widely used (Bi et al., 2024) it can be too liberal for certain datasets, potentially introducing more false positives than desirable in the context of network reconstruction (Lobo et al., 2008; Yang et al., 2015). For a more conservative approach— particularly suited to sparse networks (Poisot, 2023)—we recommend using the “prc_closest_topright” option. Alternatively, users may specify a custom *threshold (*between 0 and 1) tailored to their specific needs. While threshold selection has been amply discussed in the context of species distribution models (e.g., Liu et al., 2013) it is still not well established for network reconstruction and thus, the use of custom thresholds may allow exploring further their performance.

In practice, this option may be particularly interesting when using the “marginal_all” method, where marginal posterior probabilities ∈ [0,1]. In such cases, a threshold of 0.5 can be employed as a natural baseline: edges with *p_ij_* > 0.5 can be considered likely present (thereby identified as missing links if *D_ij_* = 0), while those with *p_ij_* < 0.5 likely absent (and potentially spurious if *D_ij_* = 1). Although this unsupervised threshold can make the identification of missing or spurious links more challenging, it offers a practical/neutral starting point. To explore plausible missing or spurious links more systematically, the top_links() function ranks node pairs by their probabilities, highlighting the most plausible candidates for further inspection (see section 2.4). For the specific task of detecting missing links, these *p_ij_* can also be used as ‘scores’ in supervised binary classification provided in *sabinaHSBM*.

The generation of the final binary network is guided by the *consistency_matrix* argument, which supports two distinct approaches for combining fold-specific predictions. The default, "average_thresholded", calculates the link mean probabilities *p_ij_* across folds and applies the mean of fold-specific thresholds for consistency. Alternatively, *sabinaHSBM* introduces a new method, "ensemble_binary", which explicitly integrates cross-validation variability into the final reconstruction. This method first applies fold-specific thresholds to the fold-specific probabilities to transform continuous matrices of each fold into binary matrices. These binary matrices are then consolidated, assigning a link as 1 if it exceeds the fold-specific threshold in the required proportion of folds. By default, this proportion is 0.1 (i.e., the new link should be in at least one fold when using 10 folds), but this proportion can be adjusted via the *ensemble_threshold* argument, with values between 0 and 1. This approach integrates fold-specific variability, ensuring that links consistently identified as significant are retained.

The *na_treatment* argument controls how undefined NA values—resulting from edges never sampled in the posterior under “marginal_all” method—are handled. By default, *sabinaHSBM* assigns such cases probability = 0 (“na_to_0”), under the assumption that rarely sampled edges are unlikely. Alternatively, you can choose “ignore_na” that exclude NAs when computing fold-averaged probabilities, thresholds, and performance metrics, effectively treating them as missing data rather than definite absences, or retaining them as NA (“keep_na”), depending on the desired treatment for undefined links.

The function evaluates model performance through fold-specific metrics, including precision, recall, sensitivity, specificity, accuracy, error rate (ERR), True Skill Statistic (TSS), and the proportion of held-out correctly recovered as 1s (the recovery link ratio, RLR). It also provides AUC values for ROC and PRC curves, along with the PRC baselines (yPRC), which represent the expected AUC-PR under random predictions.

Outputs include the reconstructed binary matrix ($new_mat), the averaged links probabilities across folds with uncertainty estimates ($res_averaged), and a detailed summary of reconstruction statistics for each fold ($stats), providing a comprehensive evaluation of the network at the fold level.

For convenience, applying the summary() function to the output of hsbm.reconstructed() returns a summary of the final reconstructed network structure—including the number of identified spurious and missing links—as well as the average of evaluation metrics across folds, providing a compact overview of overall model performance.

### 2.4 Auxiliary tools for insights and visualization

*sabinaHSBM* includes additional tools to complement the main workflow. The top_links() function ranks the most significant links based on predicted probabilities, aiding targeted exploration. The *edge_type* argument specifies the ranking type: “unrecorded” (0s likely to be 1s/missing links, ordered highest to lowest), “documented” (1s likely to be 0s/spurious links, ordered lowest to highest), while “missing” (ordered highest to lowest) and “spurious” (ordered lowest to highest) refine these rankings by focusing only on links identified as such in the final reconstructed binary matrix. To explore potential erroneous links, we recommend using the top_links() function to rank low-probability documented links (*edge_type* = "documented"). For visualizing binary bipartite matrices, the plot_interaction_matrix() function offers simple plots, accommodating both original and reconstructed networks.

For a complete description of function arguments and outputs, users can consult the help pages via the standard R documentation system (e.g., ?hsbm.input, ?hsbm.predict, ?hsbm.reconstructed, ?top_links, ?plot_interaction_matrix).

## 3 CASE STUDY: Predicting missing links in host-parasite networks

Link prediction in host-parasite networks is of crucial importance in ecology and epidemiology (Poulin, 2010), more so in a context of climate change (Morales-Castilla et al., 2021). To illustrate the functionality of *sabinaHSBM,* we use the Global Mammal Parasite Database 2.0 (Stephens et al., 2017), which compiles observed interactions between wild mammal hosts and their parasites. The dataset was filtered to include only species of the order Carnivora, and retain only records for which both hosts and parasites names were reported as valid binomial species names. For an additional analysis, we further restrict the dataset to hosts included in the phylogeny of (Faurby & Svenning, 2015). The final dataset contained 142 carnivore hosts, 695 parasites, and 2,172 documented interactions (Supplementary Data).

To structure the data for model fitting, we used the hsbm.input() function, which processes a binary interaction matrix with rows as hosts and columns as parasites. We set *n_folds* = 10 to implement 10-fold cross validation. The host-parasite interaction matrix was visualized using the plot_interaction_matrix() function (Fig. 1), revealing a sparse network with ∼2% of all possible links recorded.

**Fig. 1.**
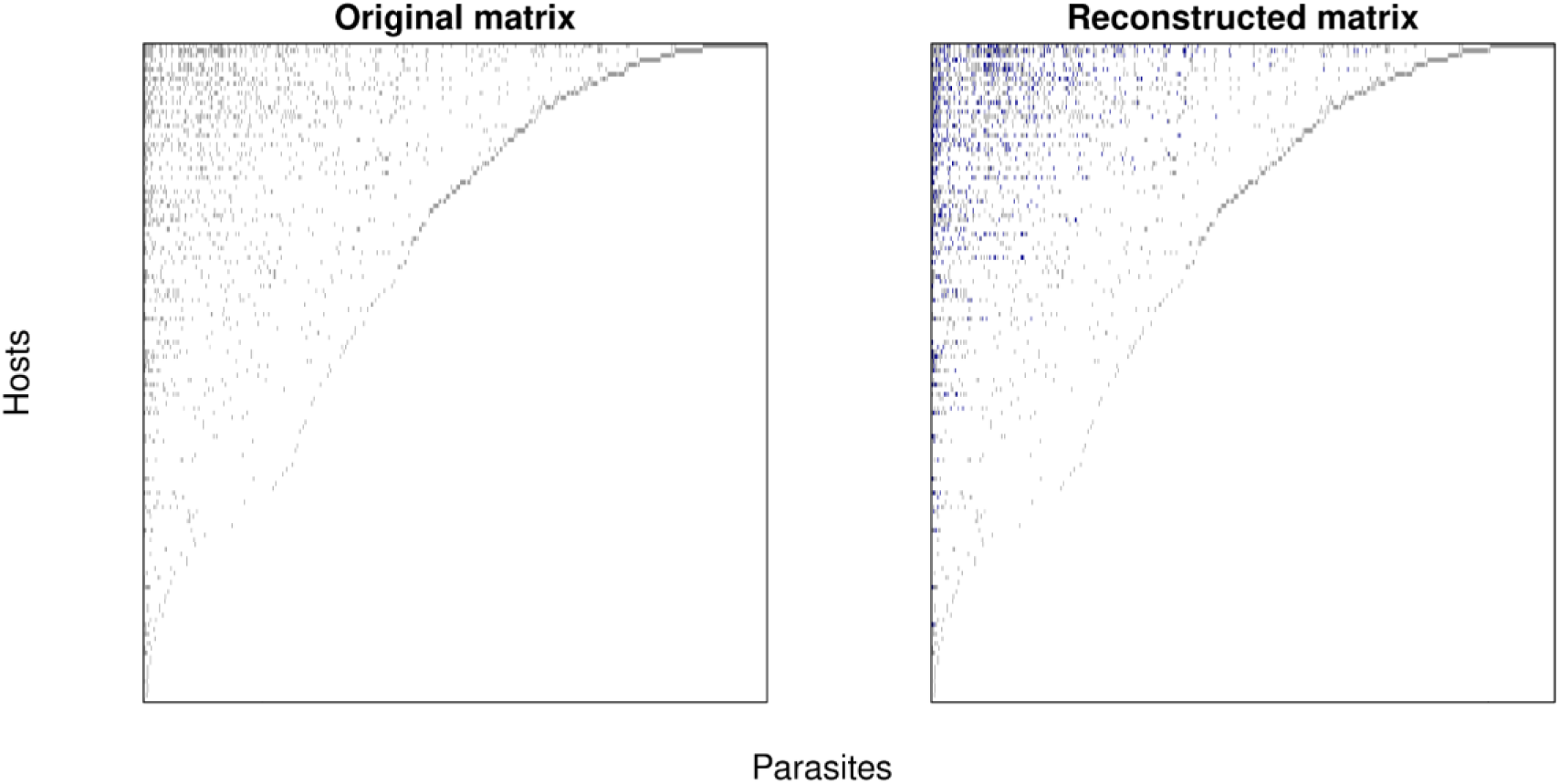
Reconstruction of the carnivore–parasite interaction matrix. Observed interaction matrix (left) and reconstructed matrix (right) obtained with *sabinaHSBM*. Rows represent carnivore hosts and columns parasite species. In the reconstructed matrix, blue cells highlight interactions predicted by the model that were absent from the empirical dataset, i.e. missing links.

To compute link probabilities, we applied the hsbm.predict() function, with the *method* = “marginal_all”, which computes marginal posterior probabilities for all node pairs. We used the default *iter* = 10,000 and *wait* = 1,000 to control the number of posterior samples and equilibration iterations, respectively. The resulting matrix of interaction probabilities provides a fully reconstructed network that can serve as a probabilistic input for Bayesian downstream analyses, allowing users to propagate uncertainty into subsequent inference steps.

To classify predicted links and evaluate network reconstruction, we used the hsbm.reconstructed() function, applying the Youden threshold from the ROC curve (*threshold* = “roc_youden”). While the “marginal_all” method outputs continuous posterior probabilities, binarizing these values serves a practical purpose: it enables interpretable, threshold-based validation and facilitates comparison with observed binary interaction data. This step allows us to compute performance metrics (e.g. AUC, RLR) and identify high-confidence missing links under a principled cutoff. The model achieved a mean AUC of 0.98 across folds and an average recovery of 39% of held-out links (RLR) (Table S1 in Supporting Information S4). The final binary matrix included 649 newly predicted interactions classified as missing links (Fig. 1), obtained using the *consistency_matrix =* “average_thresholded” option, which applies the mean of fold-specific thresholds to the averaged link probabilities. This binarized output is not intended to replace the full probability matrix but to complement it—highlighting the most robust predictions and enabling downstream comparisons that require discrete inputs.

To assess the plausibility of the most likely missing links, we used the top_links() function to extract the 10 highest-ranked undocumented interactions according to the model. A brief web literature review using Google scholar provided published evidence for 8 out of 10 of these interactions, with an additional interaction supported by reports of the parasite in closely related host species (Table S2 at Supporting Information S4).

The HSBM inferred between 3 and 4 hierarchical levels of group structure in each fold. Given that host-parasite interactions often exhibit phylogenetic signals (Davies & Pedersen, 2008) the biological relevance of these groupings was evaluated using the Mean Phylogenetic Distance (MPD) and Mean Nearest Taxon Distance (MNTD) metrics. These metrics were computed using the *picante* package (Kembel, 2010) and compared to a null model. For illustration, we present results from fold 6 (Fig. 2) although similar patterns were observed across folds (Table S5, S6). Host grouping levels for fold 6 showed phylogenetic clustering for most groupings with MPD and for some groups with MNTD, suggesting that HSBM effectively uncovered ecological and evolutionary meaningful structure in the network without prior information.

**Fig. 2.**
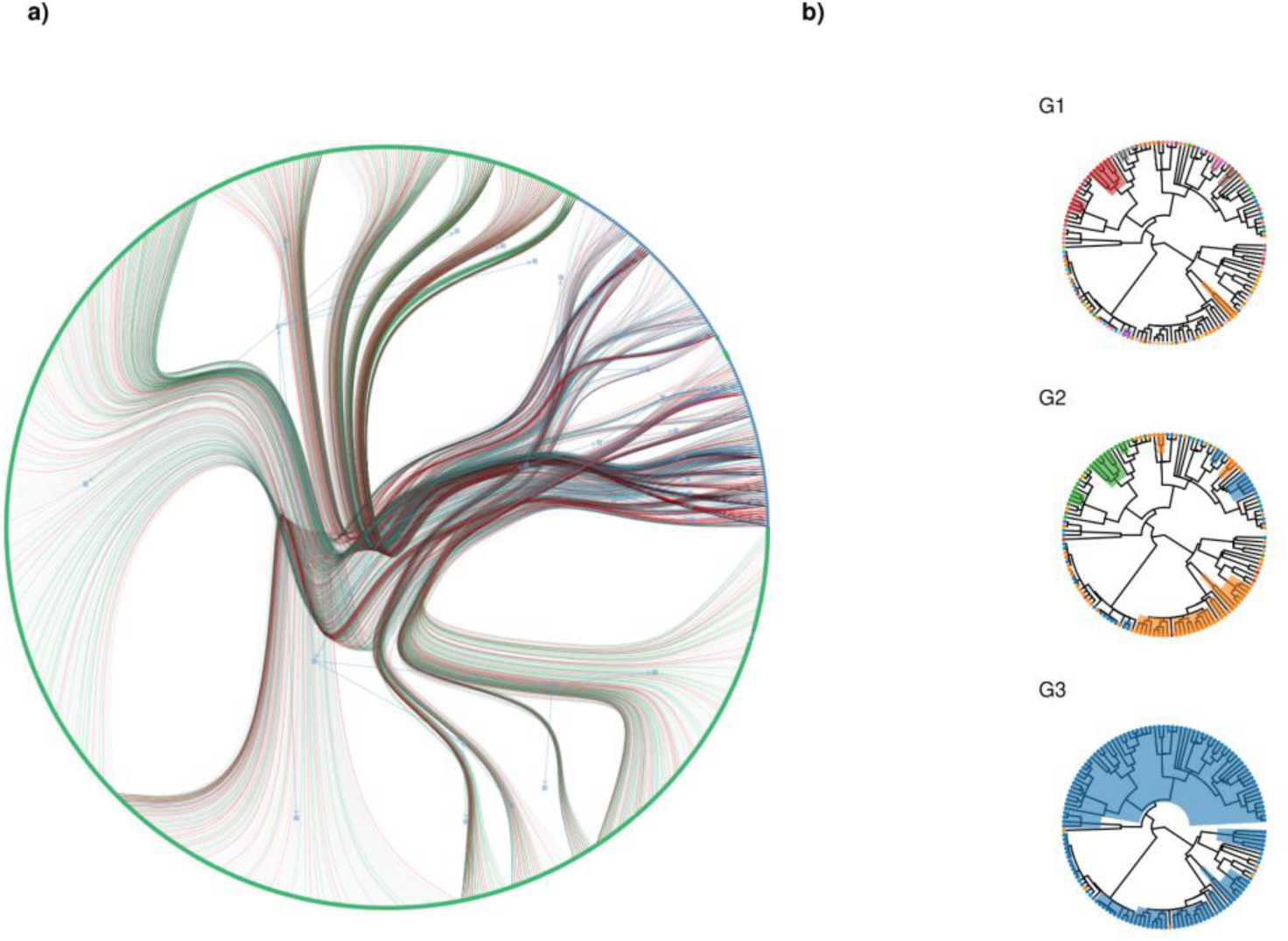
Hierarchical structure. a) Hierarchical edge bundling of the carnivore–parasite network, grouping edges by inferred blocks. b) Phylogenetic clustering of carnivore hosts at three hierarchical levels (G1–G3). Tip colors correspond to inferred groups, and clades in which all tips belong to the same group are shaded consistently to the root.

## 4 DISCUSSION

The *sabinaHSBM* package leverages the power of the hierarchical and degree-corrected stochastic block model (HSBM) for network reconstruction, offering a comprehensive and customizable set of functions that streamline the entire reconstruction workflow.

However, we emphasize that network reconstruction remains a fundamentally challenging task, and no single algorithm performs best in all contexts, as shown by Ghasemian et al. (2020) comprehensive evaluation of existing methods. Nonetheless, their results show that HSBM was the most accurate individual method in comparative analyses, performing competitively with stacked models. This was particularly evident in networks with limited data, such as many biological systems, where embedded and topological approaches performed poorly.

While being able to reconstruct networks solely from their structure would be an ideal outcome, its effectiveness depends heavily on the quality of the input data (Peixoto, 2018). In cases where datasets are small, noisy, or biased, the removal of links for cross-validation can eliminate crucial structural information, rendering reconstruction unreliable. In such situations, incorporating extrinsic data or information—such as phylogenetic relationships or species traits—can improve link prediction by supplying additional information to guide the inference process (Morales-Castilla et al., 2021). Several methods explicitly integrate such metadata, including HPprediction (Farrell et al., 2022), which uses host and parasite traits and phylogeny to predict missing interactions, or the plug-and-play approach (Dallas et al., 2017), which estimates link probabilities based on environmental or biological similarity between species pairs. However, it is important to note that doing so potentially shapes the results according to the nature and quality of the external information. In any other case, the HSBM can uncover underlying structure directly from network topology, without prior specification or the introduction of subjective decisions. This feature makes HSBM especially valuable in taxonomic groups where external data are scarce or difficult to obtain. Moreover, its hierarchical nature enables the model to capture multiple levels of structural organization, allowing for robust and reliable network reconstruction across diverse contexts.

Beyond classical ecological interaction networks, *sabinaHSBM* can be applied to a wide range of bipartite binary datasets in ecology and other disciplines where latent structure and uncertainty are relevant (see, e.g., Ghasemian et al., 2020). For instance, it can support applications in archaeology (e.g., artifact–site relationships), cultural evolution (e.g., motif– language or ritual–community associations), and historical demography or ethnography (e.g., individual–event affiliations). Beyond the humanities, *sabinaHSBM* may also prove useful in epidemiology (e.g., host–pathogen networks), user–content systems (e.g., users reviewing or tagging items in cultural or scientific repositories), or bibliometrics (e.g., authors linked to publications or journals). These broader applications illustrate the package’s versatility and its potential to support rigorous inference across diverse research domains.

## 5 CONCLUSIONS

The *sabinaHSBM* package broadens access to HSBM for ecological research and beyond. Its capabilities for link prediction, network reconstruction, uncertainty quantification, and hierarchical grouping provide a valuable tool for addressing incomplete and biased datasets, ultimately enhancing ecological and biodiversity research.

### Box1

**sabinaHSBM dual workflow for network reconstruction**

The workflow follows three steps—prepare input, predict links, and reconstruct the network—and offers two alternative strategies. The upper path illustrates a conservative "conditional_missing" mode, which recovers missing links by evaluating only unobserved interactions. The lower path shows the "marginal_air mode, which evaluates all links and can identify both missing and spurious interactions. Both strategies share the same three-step structure but differ in scope, assumptions and intended use. Functions applied in each step are indicated in the boxes

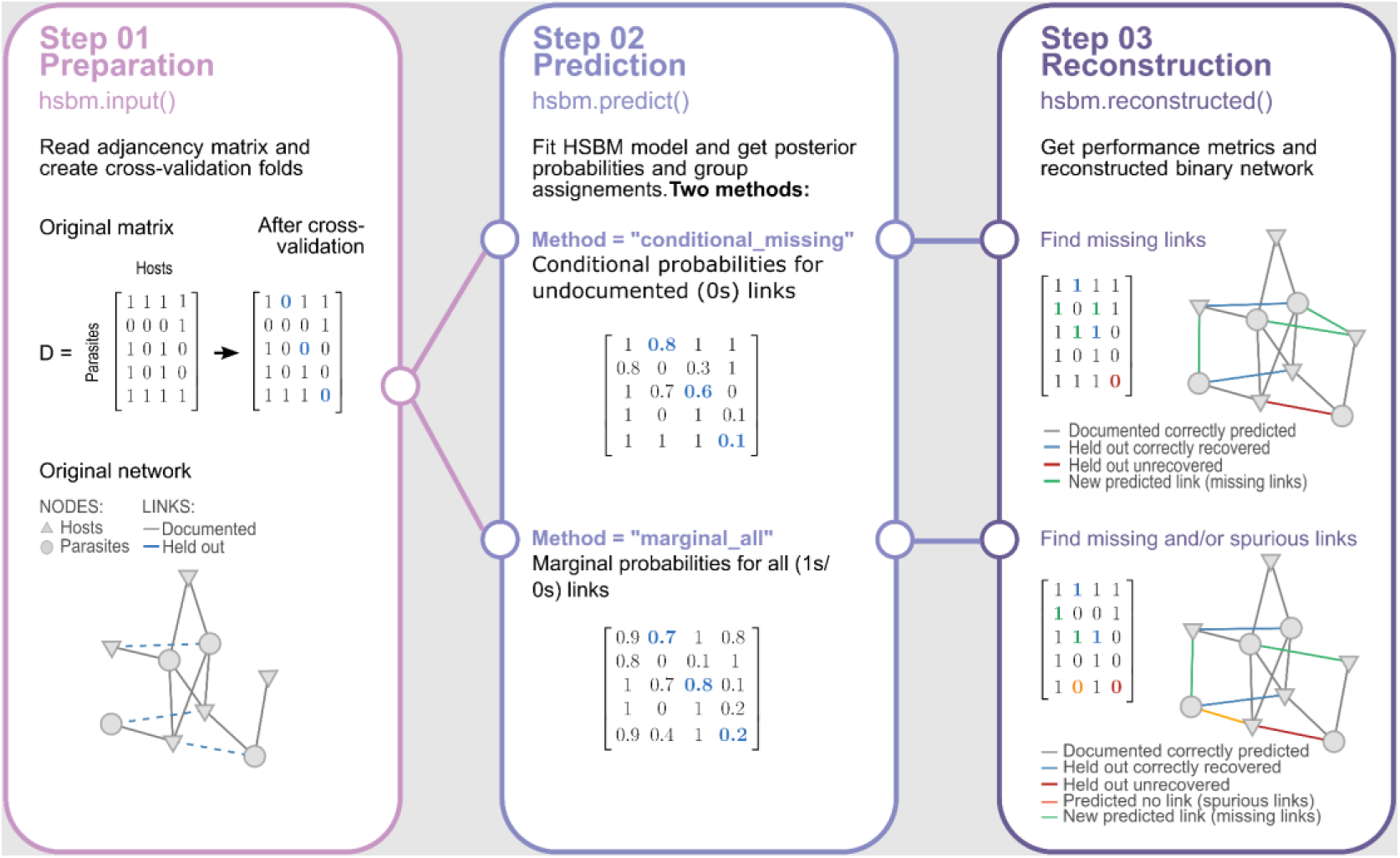

## ACKNOWLEDGEMENTS

This study was supported by the NextDive project (PID2021-124187NB-I00), funded by the Ministerio de Ciencia e Innovación (Agencia Estatal de Investigación) and the European Regional Development Fund (FEDER) — “A way of making Europe. H. L. was supported by the FPI program of the Ministerio de Ciencia e Innovacion of Spain (grant CGL2017-86926-P). I.M.-C. acknowledges funding from the Spanish Ministry for Science and Innovation (grant number PID2023-152329OB-I00 to I.M.-C.).

## AUTHOR CONTRIBUTIONS

HL and JMB developed the R package with inputs from IMC, MAR and RGM. HL and JMB jointly prepared the first version of the manuscript. All authors reviewed and contributed to the manuscript.

## DATA AVAILABILITY

Code and data are hosted in a public GitHub repository (https://github.com/h-lima/sabinaHSBM), and all materials referenced in the article will be permanently and openly accessible there.

