## Supporting Information S1 for "*sabinaHSBM*: An R package for link prediction network reconstruction using Hierarchical Stochastic Block Models"

### ***sabinaHSBM*** Docker Setup

#### Overview

This guide will help Windows users run the ***sabinaHSBM*** package within a Linux-based Docker container.

##### What's included in the Docker Image

- **Linux operating system**: Ubuntu 24.04
- **R Environment**: Pre-configured with all necessary packages, including `reticulate` for Python integration.
- **Python & graph-tool**: Python is pre-configured with `graph-tool` for network analysis.
- **sabinaHSBM**: The Docker image includes the ***sabinaHSBM*** package and all required dependencies.
- **RStudio Server**: Installed and ready to use as a browser-based R interface (user creation recommended).
- **Jupyter Notebook**: Pre-installed and ready to launch from the container for interactive coding and analysis.

##### Step-by-step setup

1. **Install Docker Desktop**

Download and install Docker Desktop for Windows from https://www.docker.com/products/docker-desktop. Registration and sign-in are required. Once Docker Desktop is installed, you can open it and run the following commands either from its integrated terminal (`>_ Terminal` button) or from the Windows Command Prompt (CMD) or PowerShell.

2. **Pull the docker image**

To download the pre-configured image from Docker Hub, run:

```
docker pull herlima/sabinahsbm
```

3. **Create and start the container**

The following command creates and runs a new Docker container named `sabinahsbm_container` using the `sabinahsbm` image. The `-p` flags are optional and map ports `8787`, `8880` from the container to the host machine, enabling access to Jupyter Notebook or RStudio Server, respectively, if needed. The `-v` flag is also optional (but recommended) and establishes a bind mount between your local directory (`/local/path/to/your/project`) and the container's bind-mounted directory (`/home/my_project`), allowing seamless file synchronization. Files created or modified in the bind-mounted directory will be directly accessible on your local machine. Alternatively, you can use a Docker volume instead of a bind mount. Unlike bind mounts, which link directly to local directory, volume are stored within a Docker filesystem.

```
docker run -it --name sabinahsbm_container \
-p 8787:8787 -p 8880:8880 \
-v "local/path/to/your/project:/home/my_project" \
sabinahsbm bash
```

After executing this command, you will enter an interactive shell, providing direct access to the container's command line.

4. **Run R, RStudio Server or Jupyter Notebook**

Once the docker container is running, you can interact with the ***sabinaHSBM*** package using different environments: through the R console, RStudio Server, or Jupyter Notebook.

- **Run R from the terminal**

  Start an R session by typing `R`. This opens an interactive console where you can load and use ***sabinaHSBM*** from the terminal environment.

- **Use RStudio Server** *(optional)*

  RStudio Server is already installed in the image.

  A default user was created with username `test` and password `sabinahsbm`. But we recommend you to set your own. Create a user and password (This step is only needed the first time) (replace `yourname` with your desired username and password):

  ```
  useradd -m yourname
  passwd yourname # you will be prompted to set a password
  ```

  Then launch RStudio Server:

  ```
  /usr/lib/rstudio-server/bin/rserver \
  --server-daemonize=0 \
  --www-port=8787 \
  --www-address=0.0.0.0
  ```

In your browser, open: http://localhost:8787

Log in with the user and password you just created, or use the default credentials.

To work in your mounted project directory `/home/my_project` use in the R console `setwd("/home/my_project"`; or go to the `Files` pane (bottom-right), click `"..."` → `"Go to folder..."` and enter `/home/my_project`.

*Note: RStudio Server runs in the foreground. Keep the terminal open while working.*

- **Use Jupyter Notebook** *(optional)*

  You can also use a Jupyter Notebook. Launch it with:

  ```
  jupyter notebook --allow-root --ip 0.0.0.0 --port=8880 --no-browser
  ```

  The terminal will display a clickable URL. Simply click the provided link or paste it into your browser to acces the Jupyter interface, ideal for coding, data visualization, and interactive analysis.

5. **Use the *sabinaHSBM* package**

Here's an example to get started:

```
setwd("/home/my_project")

# Load sabinaHSBM
library(sabinaHSBM)

# Load the data
data(dat, package = "sabinaHSBM")

# Prepare the input
myInput <- hsbm.input(
    dat,
    n_folds = 10
)

# Generate link predictions
myPred <- hsbm.predict(myInput,
                       iter = 1000,
                       method = "conditional_missing"
)

# Reconstruct the network and evaluate
myReconst <- hsbm.reconstructed(myPred,
```

```r
                                    rm_documented = TRUE,
                                    threshold = "prc_closest_topright")
summary(myReconst)

# Save the HSBM reconstructed object
#saveRDS(myReconst, file="myReconst.RData")

# Exit R
q()
```

6. **Exit the Container**

To exit the container, simply type:

```
exit
```

7. **Stop and Restart the container**

To stop the container:

```
docker stop sabinahsbm_container
```

To start the container again and access its shell, run:

```
docker start -ai sabinahsbm_container
docker exec -it sabinahsbm_container bash
```
