## Supporting Information S2 for "*sabinaHSBM*: An R package for link prediction network reconstruction using Hierarchical Stochastic Block Models"

### Using sabinaHSBM for link prediction and network reconstruction with “conditional_missing” method

This guide shows how to use document demonstrates a simple use case based on the “conditional_missing” method in *sabinaHSBM* to identify missing links (false negatives) in an incomplete network.

We use the example dataset `dat` included in the package to show key functionalities, including data preparation, link prediction, and network reconstruction.

#### Requirements and installation

There are **two ways** to use the package:

- **Option A: Docker (recommended) Non-native UNIX users** or those who prefer to avoid manual dependency installation can use the provided **ready-to-use Docker image** (includes R, Python, all dependencies, and *sabinaHSBM* pre-installed (see Supporting Information S1 for Docker container setup).

Once the container is running, simply load the package in your R session with:

```
library(sabinaHSBM)
```

- **Option B: Native UNIX installation**

**Native UNIX users** can run *sabinaHSBM* locally **if** their system includes: - R (version ≤ 4.0.4) with all required R packages (listed below) - Python ≤ with the `graph-tool` library (version ≤ 2.45)

```
# If the package is not installed, install sabinaHSBM from GitHub
 if (!requireNamespace("sabinaHSBM", quietly = TRUE)) {
   library(remotes)
   remotes::install_github("h-lima/sabinaHSBM")
 }

# Load sabinaHSBM
library(sabinaHSBM)
```

Install required R packages if not already available:

```
list.of.packages <- c(
  "dplyr",
  "parallel",
  "reshape2",
  "reticulate",
  "stringr",
  "tidyr",
  "ROCR",
  "data.table"
```

```
)

new.packages <- list.of.packages[!(list.of.packages %in% installed.packages()[, "Package"])]
if (length(new.packages) > 0) {
  install.packages(new.packages, dependencies = TRUE)
}

for (package in list.of.packages) {
  library(package, character.only = TRUE)
}
```

```
# Record starting time (optional)
start_time <- Sys.time()
```

#### Load examaple data

The dataset `dat` is a binary bipartite matrix representing a hypothetical species interactions network. Columns and rows correspond to two different types of nodes (e.g. hosts and parasites), and links (values of `1`, in red) represent interaction between them while `0` (in black) represent lack of an observed interaction. The network contains gaps, indicating potential missing links.

```
# Load the dataset
data(dat, package = "sabinaHSBM")
```

```
# Plot the simulated matrix
plot_interaction_matrix(dat, order_mat = FALSE)
```

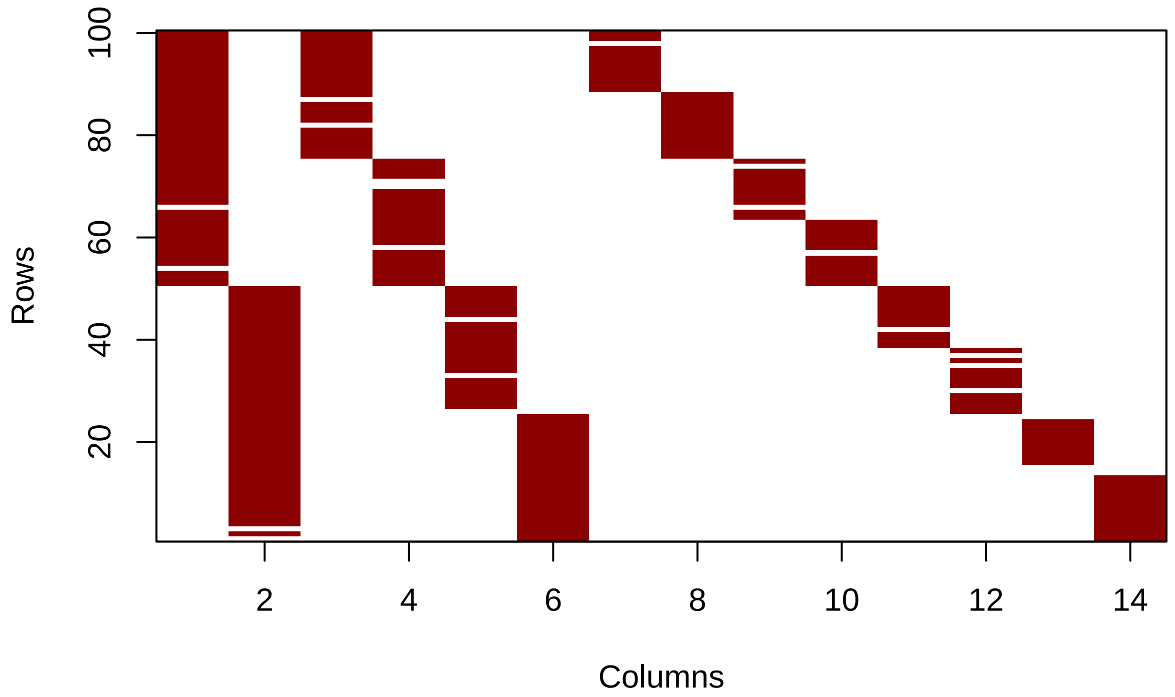


#### Preparing Input data for HSBM

The `hsbm.input` function cleans the dataset, and creates cross-validation folds by temporarily holding out subsets of observed links. Internally, it ensures that each fold masks a representative sample. In our example, it randomly splits the 277 observed links into five similar-sized subsets (because here we use 5-fold cross-validation) , holding out one subset per fold to test the model's ability to recover missing links.

```
# Prepare input object
myInput <- hsbm.input(
  dat,              # Binary bipartite matrix of observed links
  n_folds = 5       # Number of folds for cross-validation
)
```

```
# Summarizes network characteristics
summary(myInput)
```

```
##   n_rows n_cols n_links rows_single_link cols_single_link possible_links
## 1    100     14     277                1                0           1400
```

#### Predicting missing links with "conditional_missing"

The `hsbm.predict` works directly with the processed input created by `hsbm.input`. It applies HSBM to each fold's training network and computes posterior probabilities and group assignments for every possible host-parasite link. The `conditional_missing` method focuses exclusively on predicting the conditional posterior probability of unobserved links (`0s`) given the network's learned block structure, useful to identify missing links (i.e., unobserved host-parasite interactions likely to exist) in partially incomplete networks—without casting doubt on any documented interactions. Computation can be intensive, but parallelization across cores is supported.

```
# Predict missing links using HSBM
myPred <- hsbm.predict(
  myInput,          # Input data processed by hsbm.input()
  iter = 10000,      # Number of iterations ...
  wait = 1000,      # Number of iterations for MCMC equilibration
  rnd_seed = 123,      # Sets seed in python environment for reproducibility
  method = "conditional_missing",  # Prediction method
  save_blocks = TRUE,  # Save group assignments
  save_pickle = FALSE, # Save results as pickle files,
  save_plots = FALSE,  # Save hierarchical edge bundling plots
  n_cores = 2 # Number of cores to use
)
```

Predicted probabilities and group assignments are stored for each fold. Below, we extract the probabilities (p) and groups for fold 1. The lower the p, the less likely the link exists and the lower its priority; conversely, higher p values are the most promising missing interactions.

```
# View probabilities for fold 1
probabilities_fold1 <- myPred$probs[[1]]
head(probabilities_fold1)
```

```
##   v1  v2            p v1_names v2_names     edge_type
## 1  0 101 0.0010342365      sp1      SPB reconstructed
## 2  0 103 0.0027016480      sp1      SPD reconstructed
## 3  0 104 0.0004181051      sp1      SPE reconstructed
## 4  0 105 0.0004313087      sp1      SPF reconstructed
## 5  0 107 0.1574341567      sp1      SPH reconstructed
## 6  0 108 0.0008260846      sp1      SPI reconstructed
```

The group assignments provide the hierarchical clustering structure of nodes for each fold. Let's extract and examine the group assignments for fold 1:

```
# View the group/block assignments for fold 1
groups_fold1 <- myPred$groups[[1]]

# Filter one type of nodes (e.g., nodes in columns)
vnames <- colnames(myPred$data)
groups_fold1 <- groups_fold1 %>% filter(names %in% vnames)
g_cols <- grep("^G", names(groups_fold1))
groups_fold1[g_cols] <- lapply(groups_fold1[g_cols], sort)

print(groups_fold1)
```

```
##    nodes G1 G2 G3 G4 names
## 1    100  8  7  0  0   SPA
## 2    101  8  7  0  0   SPB
```

```
## 3     102  8  7  0  0   SPC
## 4     103 29  7  0  0   SPD
## 5     104 29  7  0  0   SPE
## 6     105 29  7  0  0   SPF
## 7     106 29  7  0  0   SPG
## 8     107 29  7  0  0   SPH
## 9     108 29  7  0  0   SPI
## 10    109 29  7  0  0   SPJ
## 11    110 40  7  0  0   SPK
## 12    111 72  7  0  0   SPL
## 13    112 72  7  0  0   SPM
## 14    113 72  7  0  0   SPN
```

Hierarchical group assignments provide insight into how nodes (e.g., hosts) are organized across multiple levels. At the first level (G1), nodes are divided into specific groups, reflecting fine-scale patterns. Moving to higher levels (G2, G3, G4), these groups are progressively aggregated, revealing broader patterns and relationships or communities.

### Network Reconstruction

The `hsbm.reconstructed` function generates a reconstructed binary interaction matrix by combining the probabilities across all folds and transforming them into a new binary (0/1) matrix/network from a user-specified threshold.

```
# Network reconstruction
myReconst <- hsbm.reconstructed(
  myPred,                 # Predictions processed by hsbm.predict
  rm_documented = TRUE,   # Use of documented entries during validation
  threshold = "prc_closest_topright", # Binarization threshold
  consistency_matrix = "average_thresholded" # Combine fold predictions
)
```

This output includes the averaged probability matrix, the final reconstructed matrix, and evaluation metrics. These results highlight the predicted interactions, showcasing the method's ability to detect missing links effectively. Let's explore some of the key outputs in detail.

```
# View the reconstructed network summary and evaluation metrics
summary(myReconst)
```

```
## $`Reconstructed Network Metrics`
##   obs_links unobs_links pred_links kept_links spurious_links missing_links
## 1       277        1123        281        277              0             4
##
## $`Evaluation Metrics`
##    mean_RLR  mean_auc mean_aucpr  mean_yPRC mean_prec mean_sens mean_spec
## 1 0.3835065 0.9346908  0.5134549 0.04685102 0.7588661 0.4016234 0.9926981
##    mean_ACC   mean_ERR  mean_tss
## 1 0.9650319 0.03496805 0.3943215
```

The summary provides the number of missing links, as well as key evaluation metrics, such as the recovery link ratio (RLR).

```r
# View the top potential missing links
top_links_df <- top_links(myReconst,
                          n = 10,
                          edge_type = "undocumented") # Type of edge to rank
print(top_links_df)
```

```
##    v1_names v2_names          p         sd
## 1      sp47      SPA 0.15853828 0.03890770
## 2      sp35      SPA 0.15689571 0.02573554
## 3      sp98      SPB 0.14785101 0.03562645
## 4     sp100      SPB 0.13929317 0.04113331
## 5      sp57      SPE 0.10132791 0.08325139
## 6      sp14      SPC 0.09841607 0.08716144
## 7      sp68      SPE 0.09713441 0.08583665
## 8      sp43      SPD 0.09441667 0.08877703
## 9      sp30      SPD 0.08476984 0.07569323
## 10     sp31      SPD 0.06701079 0.06251480
```

The `top-links functions` identifies the undocumented links most likely to be missing links in a network predicted by HSBM.

```r
# View the reconstructed binary matrix
plot_interaction_matrix(myReconst$new_mat, order_mat = FALSE)
```

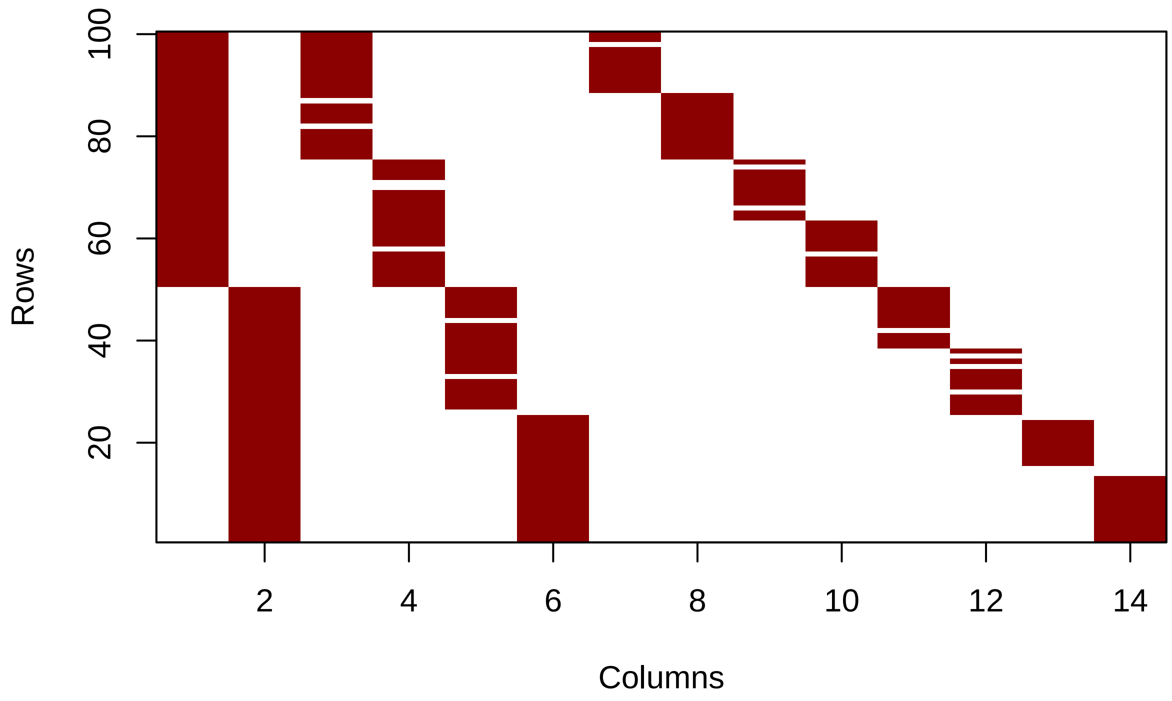

This document demonstrates the use of the *sabinaHSBM* package for network reconstruction. By applying the `conditional_missing` method, we showcased how missing links can be effectively identified and addressed in incomplete networks.

### Computing characteristics

```
# Show processing time and computer characteristics (optional)
end_time <- Sys.time()
cat("The processing time of this script took: ", end_time - start_time, "minutes\n")
```

```
## The processing time of this script took:  15.07582 minutes
```

This analysis was performed on a Dynabook with the following characteristics:

- Processor (CPU): Intel Core i7-1165G7 @ 2.80GHz (11th Gen)
- Memory (RAM): 32 GB
- Operating System: Windows 11 Pro, Version 24H2
- R Version: 4.3.3
