## Supporting Information S3 for "*sabinaHSBM*: An R package for link prediction network reconstruction using Hierarchical Stochastic Block Models"

### Using sabinaHSBM for link prediction and network reconstruction with "marginal_all" method

This guide shows how to use document demonstrates a simple use case based on the "marginal_all" method in *sabinaHSBM* to detect both missing (false negatives) and/or spurious (false positives) links in an incomplete and/or error-prone network.

We use the example dataset `dat2` included in the package to show key functionalities, including data preparation, link prediction, and network reconstruction.

#### Requirements and installation

There are **two ways** to use the package:

- **Option A: Docker (recommended) Non-native UNIX users** or those who prefer to avoid manual dependency installation can use the provided **ready-to-use Docker image** (includes R, Python, all dependencies, and *sabinaHSBM* pre-installed (see Supporting Information S1 for Docker container setup).

Once the container is running, simply load the package in your R session with:

```
library(sabinaHSBM)
```

- **Option B: Native UNIX installation**

**Native UNIX users** can run *sabinaHSBM* locally **if** their system includes: - R (version $\leq$ 4.0.4) with all required R packages (listed below) - Python $\leq$ with the `graph-tool` library (version $\leq$ 2.45)

```
# If the package is not installed, install sabinaHSBM from GitHub
 if (!requireNamespace("sabinaHSBM", quietly = TRUE)) {
   library(remotes)
   remotes::install_github("h-lima/sabinaHSBM")
 }

# Load sabinaHSBM
library(sabinaHSBM)
```

Install required R packages if not already available:

```
list.of.packages <- c(
  "dplyr",
  "parallel",
  "reshape2",
  "reticulate",
  "stringr",
  "tidyr",
  "ROCR",
```

```r
  "data.table"
)

new.packages <- list.of.packages[!(list.of.packages %in% installed.packages()[, "Package"])]
if (length(new.packages) > 0) {
  install.packages(new.packages, dependencies = TRUE)
}

for (package in list.of.packages) {
  library(package, character.only = TRUE)
}
```

```r
# Record starting time (optional)
start_time <- Sys.time()
```

#### Load example data

The dataset `dat2` is a binary bipartite matrix representing a hypothetical species interactions network. Columns and rows correspond to two different types of nodes (e.g., hosts and parasites), and links (values of `1`, in red) represent interactions between them while `0` (in black) represent lack of an observed interaction. These links are distributed in a structured pattern, with no missing information, to focus the example on identifying spurious links.

```r
# Load the dataset
data(dat2, package = "sabinaHSBM")
dat<- dat2
```

```r
# Plot the original matrix
plot_interaction_matrix(adj_mat = dat, order_mat = FALSE)
```

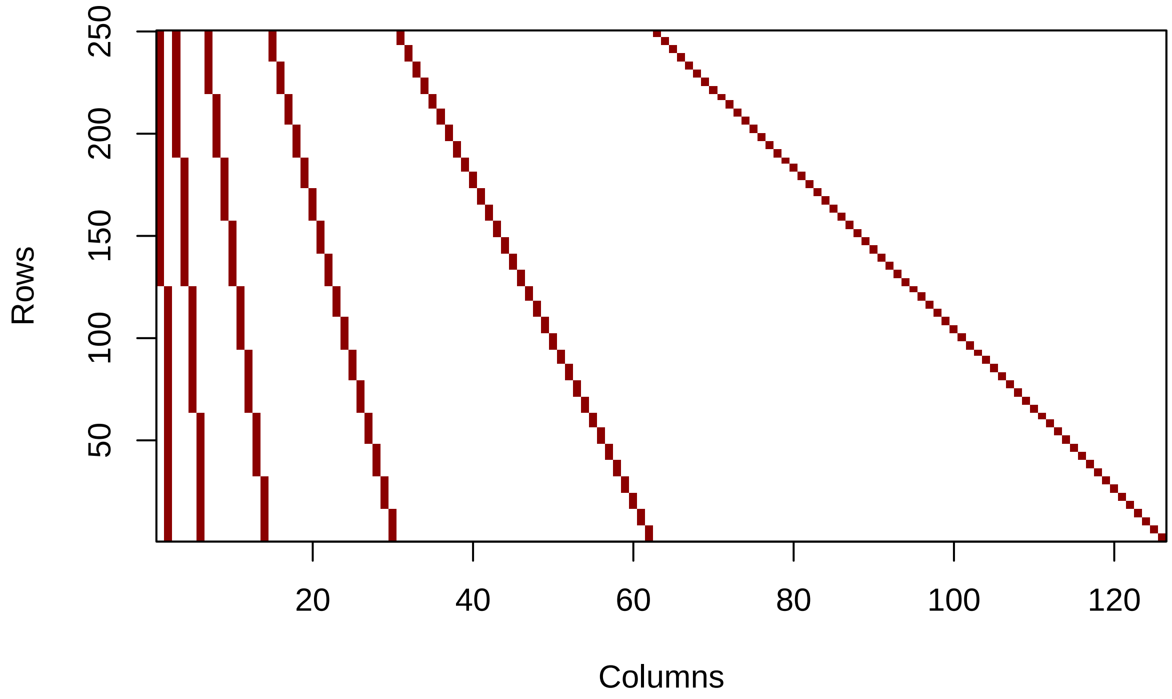


To simulate spurious links and address the `marginal_all` method, we randomly add a small number of false positives (1) to the matrix. This controlled modification allows us to demonstrate how the method can identify both spurious links (observed but potentially erroneous interactions, false positives) and missing links (unobserved but likely interactions, false negatives).

```
# Add spurious links artificially
set.seed(123)
num_spurious <- ceiling(sum(dat) * 0.01) # Proportion of spurious links to add
spurious_links <- matrix(nrow = num_spurious, ncol = 2)

for (i in 1:num_spurious) {
   repeat {
     random_row <- sample(1:nrow(dat), 1)
     random_col <- sample(1:ncol(dat), 1)

     if (dat[random_row, random_col] == 0) {
       dat[random_row, random_col] <- 1
       spurious_links[i,] <- c(random_row, random_col)
       break
     }
   }
 }
```

```
# Plot the matrix with spurious
plot_interaction_matrix(dat, order_mat = FALSE)
```

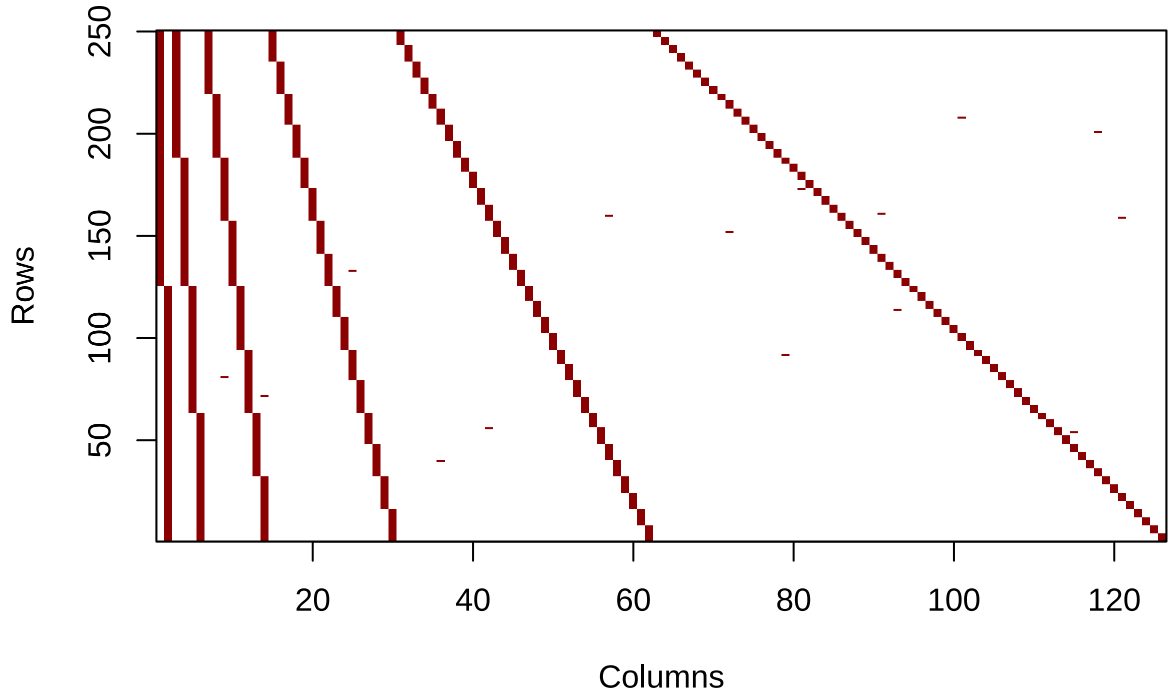


#### Prepare input data for HSBM

The `hsbm.input` function cleans the dataset and creates cross-validation folds by temporarily holding out subsets of observed links. Internally, it ensures that each fold masks a representative sample. In our example, it randomly splits all observed links into five similar-sized subsets (because here we use 5-fold cross-validation), holding out one subset per fold to test the model's ability to recover missing links.

```
# Prepare input data
n_folds <- 5       # Number of folds for cross-validation

myInput <- hsbm.input(
  dat,               # Binary bipartite matrix of observed links
  n_folds = n_folds,
  add_spurious = FALSE # Simulate spurious links
)
```

For the purposes of this example, the spurious links artificially added are labeled with the "spurious_edge" tag using the code below. This labeling allows for tracking these links throughout the analysis and network reconstruction process, although in typical use, such labeling would not be necessary.

```
myInput$edgelists <- lapply(myInput$edgelists,
                             function(x){
                                 x$x <- 1
                                 return(x)
```

```r
                             })

for(i in 1:length(myInput$edgelists)){
    el <- myInput$edgelists[[i]]
    rows <- as.numeric(el[, 1]) + 1
    cols <- as.numeric(el[, 2]) - nrow(myInput$data) + 1
    rows_cols <- paste0(rows, "_", cols)
    spurious_rows_cols <- paste0(spurious_links[, 1], "_", spurious_links[, 2])
    spurious_loc <- which(rows_cols %in% spurious_rows_cols)
    el[spurious_loc, ]$x <- 1
    el[spurious_loc, ]$edge_type <- "spurious_edge"
    myInput$edgelists[[i]] <- el
}
```

```r
# Summarizes network characteristics
summary(myInput)
```

```
##   n_rows n_cols n_links rows_single_link cols_single_link possible_links
## 1    250    126    1515                0                0          31500
```

### Predict link probabilities with "marginal_all"

The `hsbm.predict` function applies the HSBM to predict marginal posterior probabilities of all links—both observed (`1s`) and unobserved (`0s`)—so you can identify spurious and missing links in one step. The function works directly with the processed input created by `hsbm.input`.Computation can be intensive, but parallelization across cores is supported.

```r
# Define the number of cores to use
nCores <- 2

# Generate HSBM predictions
myPred <- hsbm.predict(
  myInput,                # Input data processed by hsbm.input()
  iter = 10000,           # Number of iterations
  wait= 10000,            # Number of iterations for MCMC equilibration
  rnd_seed = 123,         # Sets seed in python environment for reproducibility
  method = "marginal_all", # Method for link prediction
  save_blocks = TRUE,     # Save group assignments
  save_pickle = FALSE,    # Save results as pickle files,
  save_plots = FALSE,     # Save hierarchical edge bundling plots
  n_cores = nCores
)
```

Predicted link probabilities and group assignments are stored for each fold.

Below, we extract the link probabilities (p) for fold 1. For example, a high probability ($p = 0.91$) of an undocumented link between host 1 and parasite 2 would suggest that this interaction is likely real but was missed due to sampling limitations. Conversely, it may assign a low p (e.g., 0.12) to a recorded interaction between host and parasite, indicating it might be spurious.

```r
# View probabilities for fold 1
probabilities_fold1 <- myPred$probs[[1]]
head(probabilities_fold1)
```

```
##   v1  v2 p v1_names v2_names  edge_type
## 1  0 280 1      sp1     SPeb   held_out
## 2  0 256 1      sp1      SPg documented
## 3  0 264 1      sp1      SPo documented
## 4  0 312 1      sp1     SPkc documented
## 5  0 252 1      sp1      SPc   held_out
## 6  0 250 1      sp1      SPa documented
```

The group assignments provide the hierarchical clustering structure of nodes for each fold. Let's extract and examine the group assignments for fold 1:

```r
# View the group/block assignments for fold 1
groups_fold1 <- myPred$groups[[1]]

# Filter one type of nodes (nodes in columns)
vnames <- colnames(myPred$data)
groups_fold1 <- groups_fold1 %>% filter(names %in% vnames)
g_cols <- grep("^G", names(groups_fold1))
groups_fold1[g_cols] <- lapply(groups_fold1[g_cols], sort)

print(head(groups_fold1))
```

```
##   nodes G1 G2 G3 G4 G5 G6 names
## 1   250  5 40  6  0  2  0   SPa
## 2   251  5 40  6  0  2  0   SPb
## 3   252  5 40  6  0  2  0   SPc
## 4   253  5 40  6  0  2  0   SPd
## 5   254  5 40  6  0  2  0   SPe
## 6   255  5 40  6  0  2  0   SPf
```

Hierarchical group assignments provide insight into how nodes (e.g., hosts) are organized across multiple levels. At the first level (G1) nodes are divided into specific groups, reflecting fine-scale patterns. Moving to higher levels (G2, G3, G4) these groups (or communities) are progressively aggregated, revealing broader patterns and relationships or communities.

Next, we will explore how the model assigns specific probabilities p to different types of links ("documented," "held out," and "spurious_edge") for each fold to evaluate HSBM ability to correctly identify them. Documented links, which are real observations (true positives), should have probabilities close to 1. Held-out links, which are real observations but transformed to 0s during training for validation, should also show high probabilities. In contrast, spurious links, artificially generated (initially unobserved links transformed to 1s), should have probabilities close to 0 (unless it could be a missing link), as the model should recognize them as false links (false negatives).

```r
fold_res <- list()
for(i in 1:5){
    doc <- dplyr::filter(myPred$probs[[i]], edge_type == "documented")
    held <- dplyr::filter(myPred$probs[[i]], edge_type == "held_out")
    spur <- dplyr::filter(myPred$probs[[i]], edge_type == "spurious_edge")
```

```r
    fold_data <- c(i,
                   nrow(doc), sum(doc$p < 0.95),
                   nrow(held), sum(held$p < 0.95),
                   nrow(spur), sum(spur$p < 0.95))
    fold_res[[i]] <- fold_data
}

res_summary <- do.call(rbind, fold_res)
colnames(res_summary) <- c("Fold", "Nr doc", "Doc p <0.95",
                           "Nr held", "Held p < 0.95",
                           "Nr spur", "Spur p < 0.95")

kable(res_summary) # We use kable function from kable package to prettify output here
```

| Fold | Nr doc | Doc p <0.95 | Nr held | Held p < 0.95 | Nr spur | Spur p < 0.95 |
|---:|---:|---:|---:|---:|---:|---:|
| 1 | 1201 | 1 | 299 | 0 | 15 | 11 |
| 2 | 1200 | 0 | 300 | 0 | 15 | 7 |
| 3 | 1200 | 0 | 300 | 0 | 15 | 4 |
| 4 | 1200 | 0 | 300 | 0 | 15 | 10 |
| 5 | 1199 | 0 | 301 | 0 | 15 | 6 |

### Network Reconstruction

The `hsbm.reconstructed` function generates a reconstructed binary interaction matrix by combining predictions from all folds. The predicted matrix is transformed to binary values using a user-specified threshold.

```r
# Network reconstruction
myReconst <- hsbm.reconstructed(
  myPred,                 # Predictions processed by hsbm.predict
  rm_documented=FALSE,    # Remove documented links during validation
  na_treatment = "ignore_na", # Handle NA values in predictions
  threshold = 0.5,            # Binarization threshold
  consistency_matrix = "average_thresholded", # Combine fold predictions
  spurious_edges = TRUE,
)
```

```
## Warning in get_hsbm_results(hsbm_out, input_names = TRUE, na_treatment =
## na_treatment): Predictions obtained for 12.71% of the links. Consider
## increasing the number of iterations.
```

This output includes the final averaged probability matrix, the final reconstructed binary matrix, and evaluation metrics. These results highlight the predicted interactions, showcasing the method's ability to detect spurious and missing links effectively. Let's explore some of the key outputs.

```r
# Visualize the reconstructed binary matrix
plot_interaction_matrix(myReconst$new_mat, order_mat = FALSE)
```

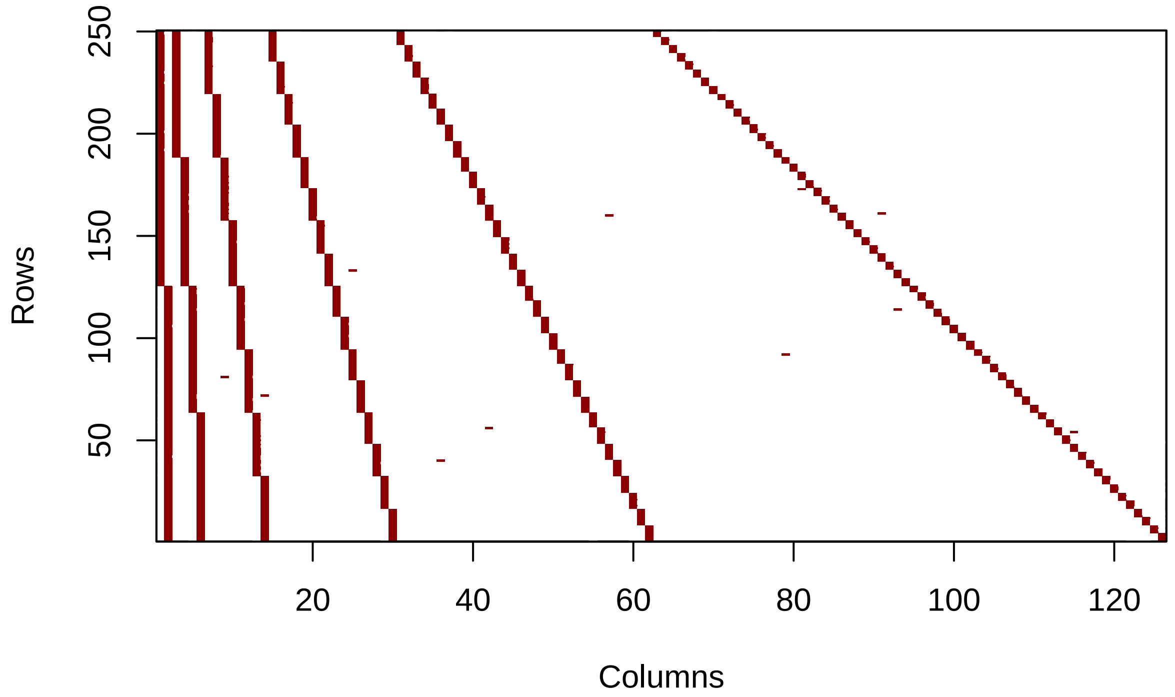


```
# View the reconstruction summary and evaluation metrics0
summary(myReconst)
```

```
## $`Reconstructed Network Metrics`
##   obs_links unobs_links pred_links kept_links spurious_links missing_links
## 1      1515       29985       1511       1511              4             0
##
## $`Evaluation Metrics`
##    mean_RLR  mean_auc mean_aucpr  mean_yPRC mean_prec mean_sens mean_spec
## 1 0.9960396 0.9990475   1.024748 0.04809524         1 0.9968317         1
##    mean_ACC    mean_ERR  mean_tss
## 1 0.9998476 0.000152381 0.9968317
```

The summary provides the number of spurious and missing links, as well as key evaluation metrics, such as the recovery link ratio (RLR).

Let's now examine the top 10 predicted links that are most likely to be spurious (false positives) by visualizing the probabilities of "documented" links.

```
# Visualize the top most likely spurious links
top_links_spurious <- top_links(myReconst,
                                n = 10,
                                edge_type = "documented")
print(top_links_spurious)
```

```
##    v1_names v2_names          p        sd
```

```
## 1      sp43     SPwd 0.02360236 0.0511687
## 2      sp92     SPqe 0.11323132 0.0495634
## 3      sp50     SPne 0.23278328 0.4157075
## 4      sp99     SPtc 0.49778978 0.3759213
## 5     sp137     SPod 0.51321132 0.4374595
## 6     sp195     SPpb 0.57315732 0.3908342
## 7      sp91     SPec 0.58407841 0.3208044
## 8     sp118      SPy 0.60238024 0.3815858
## 9     sp159     SPad 0.83748375 0.3523268
## 10    sp211     SPjb 0.86120612 0.2782042
```

This example was tailored to find spurious links and the model didn't identify any missing links at the threshold of 0.5 (but it did correctly identify 100% of the held-out links as missing). Nevertheless, we show how we can rank the most likely missing links according to the model.

```r
# Visualize the top most likely spurious links
top_links_missing <- top_links(myReconst,
                               n = 10,
                               edge_type = "undocumented")
print(top_links_missing)
```

```
##    v1_names v2_names           p          sd
## 1     sp201     SPdb 0.006200620          NA
## 2      sp45      SPg 0.005100510          NA
## 3      sp47     SPib 0.004400440 0.000000000
## 4      sp45     SPtc 0.004300430 0.000000000
## 5     sp211     SPne 0.004000400 0.000000000
## 6     sp117     SPmd 0.003900390 0.002616557
## 7     sp217     SPle 0.003700370          NA
## 8     sp216     SPab 0.003650365 0.000850085
## 9       sp7     SPsc 0.003600360          NA
## 10    sp226     SPab 0.003600360          NA
```

If the task is to do network reconstruction by inserting the most plausible missing links, we can also use the probabilities from the `marginal_all` method as 'scores' in a supervised binary classification. This is achieved by setting a discrimination threshold algorithm in `hsbm.reconstructed()` as it is done in Supporting Information S1.

This document demonstrates the use of the *sabinaHSBM* package for network reconstruction. By applying the "marginal_all" method, we showcased how spurious and missing links can be effectively identified and addressed in complex networks.

### Computing characteristics

```r
# Show processing time and computer characteristics (optional)
end_time <- Sys.time()
cat("The processing time of this script took: ",
    round(as.numeric(difftime(end_time, start_time, units = "mins", 1))), "minutes\n")
```

```
## The processing time of this script took:  107 minutes
```

This analysis was performed on a Dynabook with the following characteristics:

- Processor (CPU): Intel Core i7-1165G7 @ 2.80GHz (11th Gen)
- Memory (RAM): 32 GB
- Operating System: Windows 11 Pro, Version 24H2
- R Version: 4.3.3
