## Supporting Information S4 for "*sabinaHSBM*: An R package for link prediction network reconstruction using Hierarchical Stochastic Block Models"

Supplementary material

Herlander Lima, Jennifer Morales-Barbero, Ruben G. Mateo,
Ignacio Morales-Castilla, Miguel A. Rodríguez

### Appendix 1: MCMC sampling process in *sabinaHSBM*

The HSBM inference procedure in *sabinaHSBM* relies on the implementation in the *graph-tool* python module. It uses Markov Chain Monte Carlo (MCMC) sampling to explore the joint posterior distribution of networks $A$ and the latent block structure $b$. The process unfolds in two phases: an equilibration phase, and a posterior sampling phase.

During the equilibration phase, the model iteratively searches for convergence. In each iteration, the model performs 10 proposals to move nodes between blocks, accepting or rejecting these moves based on improvements in the posterior probability conditioned on the observed network $D$. After every iteration, the description length (DL) of the current configuration is recorded. Convergence is assessed using the `wait` argument (default: 1,000), which defines the number of successive iterations during which the minimum and maximum values of the DL must remain range stable. This period of DL stability must occur twice to ensure that convergence is not reached by chance or local fluctuations.

Once equilibration is achieved, the sampling phase begins. The model performs a number of additional iterations—defined by the `iter` argument (default: 10,000). As before, each iteration performs 10 node reassignment proposals, leading to a total of 100,000 proposals across the sampling phase. However, only the final state/configuration of each iteration is retained. These retained samples represent samples from the posterior distribution and are used to estimate link probabilities.

In the "full_reconstruction" method, marginal probabilities are computed by averaging link presence across all network configurations visited during the 10 proposals per MCMC iteration. In contrast, the "binary_classifier" method computes conditional probabilities from the final configuration of each iteration, after completing the 10 proposals. This reflects how likely the link is under each sampled block structure.

### Appendix 2: Case study results

**Table S1.** Results of reconstruction for each fold on the Carnivora dataset

| Fold | AUC | Threshold | Nr. Held-out | Pred. held-out ones | Total pred. ones |
|---|---|---|---|---|---|
| 1 | 0.99 | 0.0015 | 178 | 0.47 | 3267 |
| 2 | 0.98 | 0.0016 | 178 | 0.37 | 2775 |
| 3 | 0.99 | 0.0022 | 178 | 0.35 | 2542 |
| 4 | 0.99 | 0.002 | 178 | 0.42 | 3084 |
| 5 | 0.99 | 0.0016 | 178 | 0.45 | 3119 |
| 6 | 0.99 | 0.0024 | 179 | 0.3 | 2698 |
| 7 | 0.99 | 0.0019 | 178 | 0.33 | 2533 |
| 8 | 0.99 | 0.0019 | 178 | 0.46 | 3185 |
| 9 | 0.99 | 0.0015 | 179 | 0.37 | 2696 |
| 10 | 0.99 | 0.001 | 179 | 0.42 | 3214 |
| **Average** | **0.99** | **0.0018** | | **0.39** | **2911.3** |

**Table S2.** Literature search for most probable interactions

| Host | Parasite | Prob | SD | Evidence | Ref |
|---|---|---|---|---|---|
| Neovison vison | Mesocestoides lineatus | 0.19 | 0.27 | Confirmed | [1] |
| Lynx rufus | Carnivore protoparvovirus 1 | 0.12 | 0.31 | Confirmed | [2] |
| Procyon lotor | Yersinia pestis | 0.08 | 0.08 | Confirmed | [3] |
| Meles meles | Eucoleus aerophilus | 0.06 | 0.06 | Confirmed | [4] |
| Lynx rufus | Felid alphaherpesvirus 1 | 0.06 | 0.04 | Plausible. Artificial innoculation resulted in asymptomatic infection. | [5] |
| Procyon lotor | Ctenocephalides felis | 0.04 | 0.07 | Confirmed | [6] |
| Nyctereutes procyonoides | Capillaria aerophila | 0.03 | 0.03 | Confirmed for parasite synonym name Eucoleus aerophilus | [7] |
| Lutra lutra | Molineus patens | 0.03 | 0.03 | Confirmed | [8] |
| Nyctereutes procyonoides | Toxoplasma gondii | 0.03 | 0.02 | Confirmed | [9] |
| Neovison vison | Macracanthorhynchus catulinus | 0.03 | 0.02 | No evidence found. Infection of other members of Mustela genus. | [10] |

**Table S3.** Mean phylogenetic distance for grouping levels found for hosts during Hierarchical Stochastic Block Model inference for fold 6 on the Carnivora dataset. Results shown for groups with at least 10 hosts.

| Level | Nr hosts | Obs. | Null mean | Null sd | p | p<0.05 |
|---|---|---|---|---|---|---|
| **Level 1 (nr groups: 12)** | | | | | | |
| | 32 | 102.419 | 101.197 | 2.166 | 0.669 | |
| | 19 | 61.710 | 101.346 | 3.175 | 0.001 | * |
| | 22 | 103.088 | 101.360 | 2.818 | 0.704 | |
| | 10 | 82.213 | 101.422 | 5.928 | 0.014 | * |
| | 15 | 88.617 | 101.362 | 3.988 | 0.011 | * |
| **Level 2 (nr groups: 4)** | | | | | | |
| | 69 | 99.232 | 101.318 | 1.058 | 0.040 | * |
| | 27 | 61.427 | 101.371 | 2.351 | 0.001 | * |
| | 39 | 83.234 | 101.240 | 1.824 | 0.001 | * |
| **Level 3 (nr groups: 2)** | | | | | | |
| | 135 | 100.806 | 101.258 | 0.194 | 0.019 | * |
| | 135 | 100.806 | 101.273 | 0.188 | 0.012 | * |

**Table S4.** Mean nearest taxon distance for grouping levels found for hosts during Hierarchical Stochastic Block Model inference for fold 6 on the Carnivora dataset. Results shown for groups with at least 10 hosts.

| Level | Nr hosts | Obs | Null mean | Null sd | p | p<0.05 |
|---|---|---|---|---|---|---|
| **Level 1 (nr groups: 12)** | | | | | | |
| | 32 | 26.387 | 25.356 | 3.119 | 0.623 | |
| | 19 | 23.968 | 31.671 | 5.382 | 0.076 | |
| | 22 | 28.264 | 29.647 | 4.274 | 0.372 | |
| | 10 | 22.180 | 44.657 | 10.077 | 0.014 | * |
| | 15 | 25.960 | 35.553 | 6.664 | 0.066 | |
| **Level 2 (nr groups: 4)** | | | | | | |
| | 69 | 19.928 | 18.884 | 1.347 | 0.787 | |
| | 27 | 22.459 | 27.296 | 3.615 | 0.094 | |
| | 39 | 15.087 | 23.578 | 2.579 | 0.002 | * |
| **Level 3 (nr groups: 2)** | | | | | | |
| | 135 | 15.216 | 15.358 | 0.262 | 0.288 | |
| | 135 | 15.216 | 15.354 | 0.262 | 0.279 | |

**Table S5.** Mean phylogenetic distance of inferred groupings compared to null model. G1, G2, G3, G4 are the grouping levels. In each level the numbers refer to how many inferred grouping were found to be significant ($p < 0.05$) compared to a null model, over all groupings with more than 10 taxa. So 2/4 means that two inferred groupings were found to be significantly clustered on a total of 4 inferred groupings with more than 10 taxa.

| Fold | G1 | G2 | G3 | G4 |
|---|---|---|---|---|
| 1 | 3/6 | 2/3 | 1/1 | 1/1 |
| 2 | 2/5 | 2/3 | 0/1 | NA |
| 3 | 2/4 | 2/4 | 0/1 | 0/1 |
| 4 | 2/4 | 1/3 | 0/1 | NA |
| 5 | 2/4 | 4/4 | 1/1 | NA |
| 6 | 3/5 | 2/3 | 1/1 | 1/1 |
| 7 | 4/6 | 2/3 | 0/1 | NA |
| 8 | 2/5 | 3/3 | 0/1 | NA |
| 9 | 2/4 | 2/3 | 0/1 | NA |
| 10 | 3/5 | 4/4 | 0/1 | NA |
| **Average** | **0.52** | **0.72** | **0.3** | **0.67** |

**Table S6.** Mean nearest taxon distance of inferred groupings compared to null model. G1, G2, G3, G4 are the grouping levels. In each level the numbers refer to how many inferred grouping were found to be significant ($p < 0.05$) compared to a null model, over all groupings with more than 10 taxa. So 2/4 means that two inferred groupings were found to be significantly clustered on a total of 4 inferred groupings with more than 10 taxa.

| Fold | G1 | G2 | G3 | G4 |
|---|---|---|---|---|
| 1 | 0/6 | 1/3 | 0/1 | 0/1 |
| 2 | 0/5 | 0/3 | 0/1 | NA |
| 3 | 0/4 | 1/4 | 0/1 | 0/1 |
| 4 | 0/4 | 0/3 | 1/1 | NA |
| 5 | 1/4 | 0/4 | 0/1 | NA |
| 6 | 1/5 | 1/3 | 0/1 | 0/1 |
| 7 | 2/6 | 1/3 | 0/1 | NA |
| 8 | 1/5 | 2/3 | 1/1 | NA |
| 9 | 0/4 | 0/3 | 0/1 | NA |
| 10 | 0/5 | 1/4 | 0/1 | NA |
| **Average** | **0.1** | **0.22** | **0.2** | **0** |
